# Early Life Outcomes of Prenatal Exposure to Alcohol and Synthetic Cannabinoids in Mice

**DOI:** 10.1101/2025.01.27.635118

**Authors:** Siara K. Rouzer, McKay Domen, Aisley George, Abigail Bowring, Rajesh C. Miranda

## Abstract

This study explores the effects of prenatal co-exposure to alcohol and synthetic cannabinoids on offspring viability, physical development, and neurobehavioral outcomes in young adulthood. The aim is to identify distinct outcomes of co-exposure compared to single-drug exposures and to examine potential sex-specific vulnerabilities in motor coordination and exploratory behaviors. Pregnant C57Bl/6J mice were assigned to one of four treatment groups: Control, Alcohol-exposed, Cannabinoid-exposed, or Alcohol+Cannabinoid-exposed, with drug administration occurring between Gestational Days 12–15. Offspring were first evaluated at birth for survival, physical malformations, and developmental delays. Subsequently, young adult offspring were assessed for motor coordination using rotarod tests and exploratory behavior using open field tests. Our results indicate that alcohol and cannabinoid co-exposure significantly reduced offspring survival and litter sizes compared to controls. Non-viable offspring displayed craniofacial abnormalities, limb malformations, and developmental delays. Behavioral assessments in young adulthood demonstrated that all forms of prenatal drug exposure impaired motor coordination in males, while alcohol and cannabinoid exposures independently produced impairments in females. In the open field test, co-exposed male offspring exhibited reduced center exploration, indicative of anxiety-like behavior. Co-exposed offspring, regardless of sex, demonstrated hyperactivity, characterized by increased speed and distance traveled. Together, these findings underscore the heightened risks associated with prenatal polysubstance exposure, which exacerbates offspring mortality and induces sex-specific neurobehavioral deficits. This study highlights the distinct outcomes associated with prenatal co-exposure, and the need for future research to investigate underlying mechanisms driving these developmental disruptions and sex-specific susceptibilities.

**Structured Abstract:** *Purpose:* This study investigates the effects of prenatal co-exposure to alcohol and synthetic cannabinoids on offspring viability, physical development, and neurobehavioral outcomes in young adulthood. The goal of this investigation is to determine whether prenatal co-exposure produces distinct outcomes from single-drug exposures, including sex-specific vulnerabilities in motor coordination and exploratory behaviors.

*Methods:* Pregnant C57Bl/6J mice were randomly assigned to one of four treatment groups: drug-free controls, alcohol (ALC)-exposed, cannabinoid (CP-55,940, CB)-exposed or ALC+CB-exposed, with drug exposure occurring between Gestational Days 12-15. Offspring viability, physical malformations, and developmental delays were first assessed at birth. Then, behavioral evaluations, including rotarod and open field tests, were conducted on young adult offspring (Postnatal Days 100–120).

*Results:* ALC+CB exposure significantly decreased litter survival (*p* = 0.006) and offspring viability compared to controls. Non-viable offspring exhibited craniofacial abnormalities, limb malformations, and developmental delays. Assessments of rotarod performance revealed that all exposures reduced motor coordination in males compared to controls (*p* < 0.05), while ALC and CB exposures alone produced this outcome in females. Open field tests indicated that ALC+CB exposure reduced time in the center of the arena in male offspring exclusively, while this same exposure increased hyperactivity compared to single-drug and control groups, independent of sex (*p* < 0.05).

*Conclusions:* Prenatal co-exposure to alcohol and synthetic cannabinoids exacerbates offspring mortality and induces sex-specific deficits in neurobehavioral motor outcomes. These findings highlight the distinct risks of polysubstance exposure during pregnancy and underscore the need for targeted interventions to mitigate the effects of prenatal polysubstance exposure on offspring health outcomes.

**Highlights:**

- Alcohol and cannabinoid co-exposure significantly reduces offspring survival.
- Reduced survival is associated with offspring craniofacial and limb malformations.
- Prenatal exposures produce sex-specific impairments in motor coordination.
- Co-exposed offspring display hyperactivity in young adulthood.

## Introduction

Alcohol and cannabis are two of the most used substances by individuals of child-bearing age (Rouzer et al., 2023; Subbaraman & Kerr, 2015), and next to nicotine, are the most frequently co-used substances in human populations (Singh, 2019). Moreover, alcohol and cannabinoid co-users are most likely to be young adults of reproductive age (Subbaraman & Kerr, 2015), the age group that is also most likely to experience unplanned pregnancy (Brown & Eisenberg, 1995).

However, rates of prenatal alcohol and cannabinoid co-exposure are difficult to quantify, due to the societal stigma surrounding alcohol and cannabis use by pregnant individuals (Corrigan et al., 2017; Raifman et al., 2024). Conservative estimates place the prevalence of prenatal alcohol exposure (PAE) at ∼10% (Popova et al., 2017), while prenatal cannabis exposure (PCE) is reported among 3-35% of pregnancies (Nashed et al., 2021). Given that half of young adult cannabis users also engage in alcohol co-use (Yurasek et al., 2017), alcohol and cannabinoid co-exposure is likely a prevalent form of prenatal exposure.

Importantly, independent exposures to either alcohol or cannabinoids have been associated with compromised fetal and maternal health at the time of delivery. In humans, confirmed exposure to either substance has been associated with increased risk for adverse neonatal outcomes and pregnancy complications, including pre-term delivery and infants born small for gestational age (Addila et al., 2021; Ayonrinde et al., 2021; B. A. Bailey & R. J. Sokol, 2011; G. Bandoli et al., 2023; Petrangelo et al., 2019; Popova et al., 2021). However, reports of prenatal and early-postnatal complications associated with individual exposures are not consistent across all clinical assessments (as previously described in (Rouzer et al., 2024)), indicating that the relationship between prenatal drug exposure and offspring health risk is likely moderated by other contributing factors. Given the prevalence of co-use of alcohol and cannabinoids among young adults, we propose that one of these moderating factors may be the influence of *prenatal polysubstance use*.

At present, there is minimal research comparing offspring outcomes from alcohol and cannabinoid co-exposures to single-drug exposures. Within this under-investigated research area are several independent studies demonstrating that co-exposure can produce distinct and sex-specific outcomes in offspring, affecting fetal intrauterine blood flow (Rouzer et al., 2024), viability at birth and birth weight, behavioral and cognitive performance, and underlying neurological systems, including those associated with memory, endocannabinoid signaling, and the *Sonic Hedgehog* gene (reviewed in detail in (Rouzer et al., 2023)). However, select studies have also reported that co-exposure does not always augment outcomes from singular exposures, instead mimicking the outcomes of individual exposures (Ornelas et al., 2024; Reid et al., 2024). These conflicting reports further reflect the need to investigate and identify the contexts under which combined exposure may increase risk for offspring neurodevelopmental impairments throughout the lifespan.

Our study appends this limited body of research using gestational exposures to alcohol and/or synthetic cannabinoid CP-55940 in mice, controlling for the timing and quantities of drug exposures across pregnancies. Following characterization of birth outcomes, offspring aged into young adulthood and were assessed for motor deficits, a well-characterized consequence of offspring exposure to alcohol (Bhatara et al., 2006; Breit et al., 2019a; Cheng et al., 2018; Connor et al., 2006; Lucas et al., 2014; Mick et al., 2002) or cannabinoids (Breit et al., 2019b; Campolongo et al., 2011; de Salas-Quiroga et al., 2015; Hussain et al., 2022; A. F. Scheyer et al., 2019; Shabani et al., 2011). Our investigation reveals prenatal cannabinoid-driven effects, including the loss of offspring viability, which is associated with the expression of craniofacial malformations, underdeveloped limbs and digits, and immaturity for gestational age. In early adulthood, prenatal co-exposure reduces motor coordination and time spent in the center of an open field seen only in male offspring, while increases in hyperactivity were sex-independent in this group. Notably, all exposures produce deficits in motor behaviors, highlighting that this developmental domain is commonly targeted by intrauterine alcohol and cannabinoid exposures.

## Methods

### Generation of prenatally-exposed offspring

All breeding and prenatal drug exposure procedures are described in detail in our previous publication using this co-exposure model (Rouzer et al., 2024) and were performed with the oversight and approval of Texas A&M University’s Institutional Animal Care Committee. Briefly, breeding protocols involved overnight co-housing of one male with two female C57Bl/6J mice (Jackson Laboratories, ME, Strain #000664) and successful mating was confirmed by the presence of a sperm plug (Gestational Day 1). Following pregnancy confirmation, females were assigned to one of four groups: **control (CON)**, **alcohol-only (ALC)**, **cannabinoid-only (CB)**, or **both substances (ALC + CB)**. From Gestational Days (G) 12 to 15, females received daily intraperitoneal injections of either CP-55,940 (750 µg/kg in 10% DMSO/saline, Tocris Bioscience) or volume-equivalent 10% DMSO/saline. This was followed by a 30-minute exposure to either ethanol or vapor (95% ethanol) or room air using a passive e-vape system (La Jolla Alcohol Research Inc., La Jolla, CA). Dams were visually assessed daily to verify their well-being during the prenatal exposure period leading up to delivery. Notably, any pregnant dams that exhibited physical distress or experienced delivery complications were euthanized according to approved institutional protocols (IACUC 2021-0331).

### Parameters for characterizing deceased offspring features

As previously described in our prenatal characterization of this co-exposure model, we observed instances of maternal and fetal fatality following, but not preceding, drug exposures (Rouzer et al., 2024), particularly among CB and ALC+CB litters. To investigate the effects of prenatal drug exposure on periconceptional offspring mortality, six non-viable litters were analyzed following euthanasia of the dam according to institutional protocol. Deceased offspring were photographed and these were independently reviewed by four researchers, who generated individual assessments of physical abnormalities and quantified resorptions. Analyses were conducted using defined parameters across multiple anatomical and developmental categories (described in Table 1), and ratings of deceased offspring physical malformations were reviewed for inter-rater reliability using intraclass correlation coefficient (ICC) analysis. Developmental delays were assessed using four criteria: skull shape, snout shape, pinna placement, and digit formation. Each feature was assigned to a Theiler stage based on visual appearance (Richardson et al., 2013), independent of gestational age at dissection. The discrepancy between the estimated Theiler stage and the actual Theiler stage was calculated as follows: **Difference** = Estimated Theiler stage – Actual Theiler stage. For example, if the fetal snout was estimated at Theiler Stage 22, but dissected at Theiler Stage 23, it would receive a score of -1.

**Table 1.**
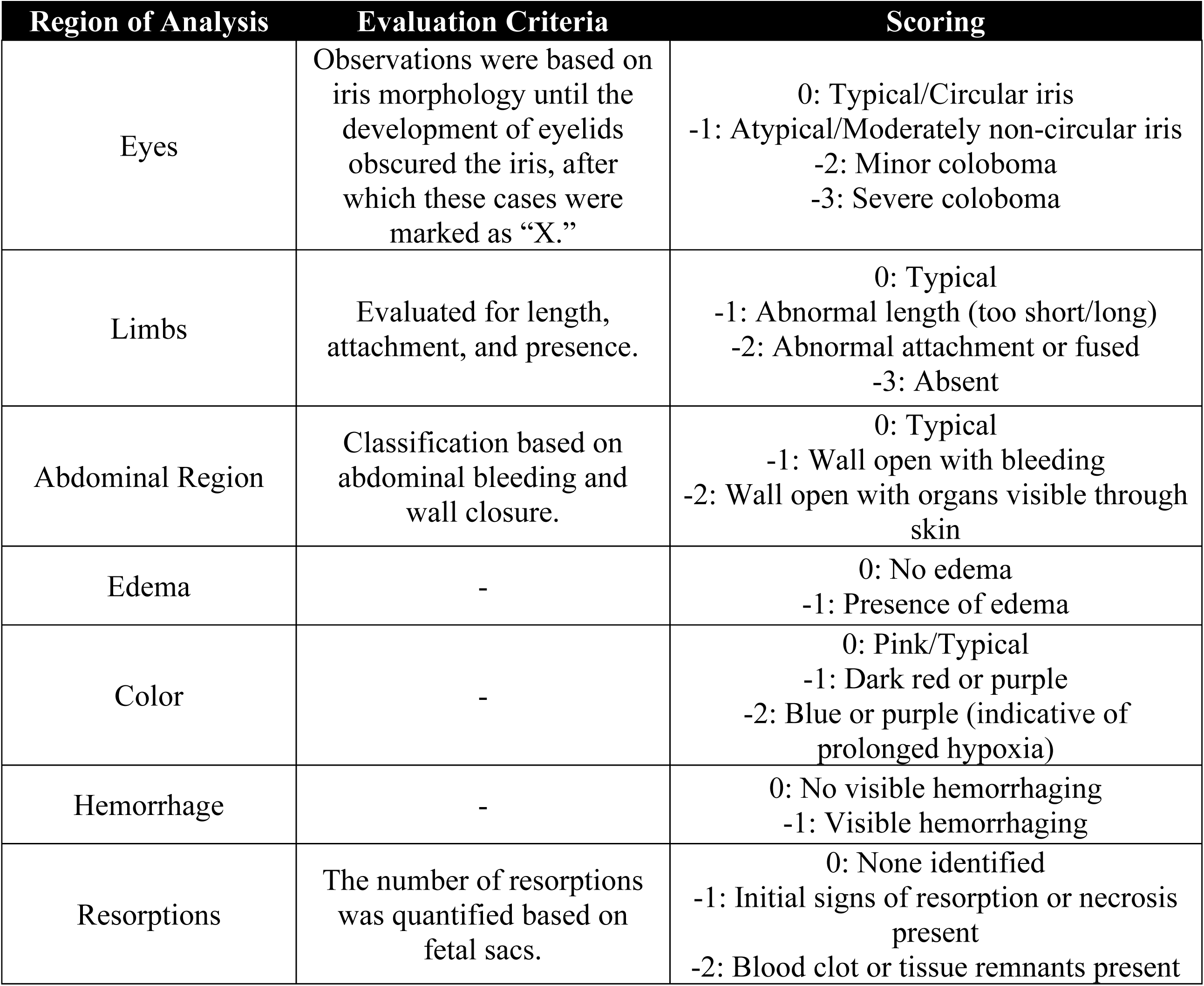
Evaluation criteria for physical assessments of non-viable offspring.

#### Data Processing and Reporting

After individual scoring, numerical data were averaged across reviewers and compiled into a single summary table. If a value could not be confidently determined from an image or fetus, reviewers provided an “X” for that metric. Variables marked as “X” by more than one reviewer were excluded from numerical scoring and retained as “X” in the final dataset. Due to the selective impact of exposures on litter viability, only certain exposure groups were represented, and no statistical analyses were performed.

### Assessments of offspring Rotarod coordination

In young adulthood (Postnatal Day [PD]100-120), male and female offspring from all exposure groups underwent rotarod testing (Rotarod Series 8, IITC Life Science Inc.) over three consecutive days. Each day, animals acclimated for 10 min in a red-light illuminated behavioral testing room, where all sessions were conducted under red-light conditions. On Day 1, animals completed a Habituation task, in which they were required to balance on the rotarod at 5 rotations-per-minute (RPM) for 120 sec. Animals failing to meet this threshold were given a 10 min break before undergoing up to two additional attempts. The number of attempts required to meet this threshold was recorded for each subject, and successful subjects advanced to subsequent testing days, which employed a new testing protocol. Each testing day was identical, and consisted of three trials with a 10 min inter-trial interval. Immediately after placement on the Rotarod, rod speed increased from 4 to 40 RPM over 108 sec. If subjects fell, their time to fall was recorded, and subjects were returned to their homecage for an inter-trial resting period. If subjects remained balanced on the rod for 150 seconds, demonstrating 30+ seconds of balance at the top speed, the trial was terminated, the maximum time of 150 seconds was recorded, and the subject was returned to their homecage. All subjects had *ad libitum* access to food and water in their homecages throughout the testing period.

### Assessments of offspring Open Field activity

Subjects were evaluated for locomotor activity and exploratory behavior using an open field test in a 50 cm x 50 cm arena with 50 cm high walls, under dim lighting conditions. After 10min to acclimate to the mouse behavioral testing room, each subject was placed at the center of the arena and allowed to explore freely for 10 minutes. An overhead camera connected to EthoVision XT software (Noldus Information Technology, The Netherlands) automatically recorded and analyzed subject movements, and provided heatmaps of subject activity based on designated exposure/sex. The software tracked and provided numerical outputs for 1) time spent in the center of the open field arena, 2) times entering the center from the periphery, 3) total distance travelled during testing, and 4) average speed travelled during testing.

### Statistics

Litter mortality was analyzed using a between-subjects analysis of variance (ANOVA), with independent variables of ALC exposure (Yes/No) and CB exposure (Yes/No). Offspring weights were analyzed using a mixed-effects ANOVA with independent variables of ALC exposure, CB exposure, and Age. Assessments of physical malformations in deceased offspring were reviewed for inter-rater reliability using Interclass correlation (ICC) analysis, with ICC estimates and their 95% confident intervals calculated based on a mean-rating (*k* = 4), absolute-agreement, 2-way random-effects model. Sex differences in control offspring performance on the rotarod were assessed using a mixed-effects ANOVA with independent variables of Sex (Male/Female), Testing Day (Day 1/Day 2), and Trial # within day (Trial 1/Trial 2/Trial 3). Sex differences in control offspring metrics in the Open Field were assessed using unpaired t-tests, comparing males and females. The effects of prenatal exposure on days to habituate to the Rotarod task, as well as Open Field measures of Time in Center, Times Entering Center, Speed Travelled, and Distance Travelled, were analyzed using a between-subjects ANOVA with independent variables of ALC exposure, CB exposure, and Sex. Within each sex, the effects of prenatal exposure on rotarod performance were assessed using a mixed-effects ANOVA with independent variables of prenatal exposure (CON, ALC, CB, or ALC+CB), Testing Day, and Trial #. Finally, within each sex, the effects of drug exposure on first and last Rotarod trials were assessed using a mixed-effects ANOVA with independent variables of ALC exposure, CB exposure, and Trial (First/Last). In all statistical assessments where sex was not a statistically significant main effect or interacting variable, data were collapsed to perform follow-up assessments investigating sex-independent effects of prenatal exposures. For all assessments, in the event of a significant main effect or interaction between independent variables, post-hoc Tukey tests were used to determine significant differences between individual groups. All data were assessed for outliers using the ROUT method of regression with a false-discovery rate of 1%, and identified outliers were removed from statistical analyses. Group differences were considered significant at p ≤ 0.05. ICC analysis was performed using SPSS (v28, IBM, NY). All other statistical tests were performed using Prism (v9, GraphPad Software, MA).

## Results

### Prenatal polysubstance exposure decreases offspring viability and litter sizes

In postnatal characterization of our ALC and CB co-exposure model, we found a significant main effect of CB exposure on litter survival [*F*(1, 36) = 9.559, *p* = 0.004] and a non-significant main effect of ALC exposure on litter survival [*F*(1, 36) = 3.204, *p* = 0.082]. Post-hoc analyses revealed that this effect was driven by the ALC+CB group, which demonstrated significantly lower litter survival (∼33%) than the control group (*p* = 0.006; Supplementary Table 1; Figure 1A). Furthermore, among viable litters, CB exposure significantly impacted the number of live pups/birth [*F*(1, 20) = 13.88, *p* = 0.001], with ALC+CB litters bearing fewer offspring than control litters (*p* = 0.067) and ALC litters (*p* = 0.027; Supplementary Table 2; Figure 1B). Among surviving offspring, there was a trend, though statistically non-significant, for an interaction between ALC and CB exposure on litter sex ratios [*F*(1, 19) = 3.706, *p* = 0.069], with no significant post-hoc comparisons (Supplementary Table 3; Figure 1C). It is worth noting that the lack of statistical significance may reflect an insufficient sample size due to the significant loss of offspring viability associated with ALC+CB exposure, and given that twice as many male offspring survived among these litters as female offspring.

**Figure 1.**
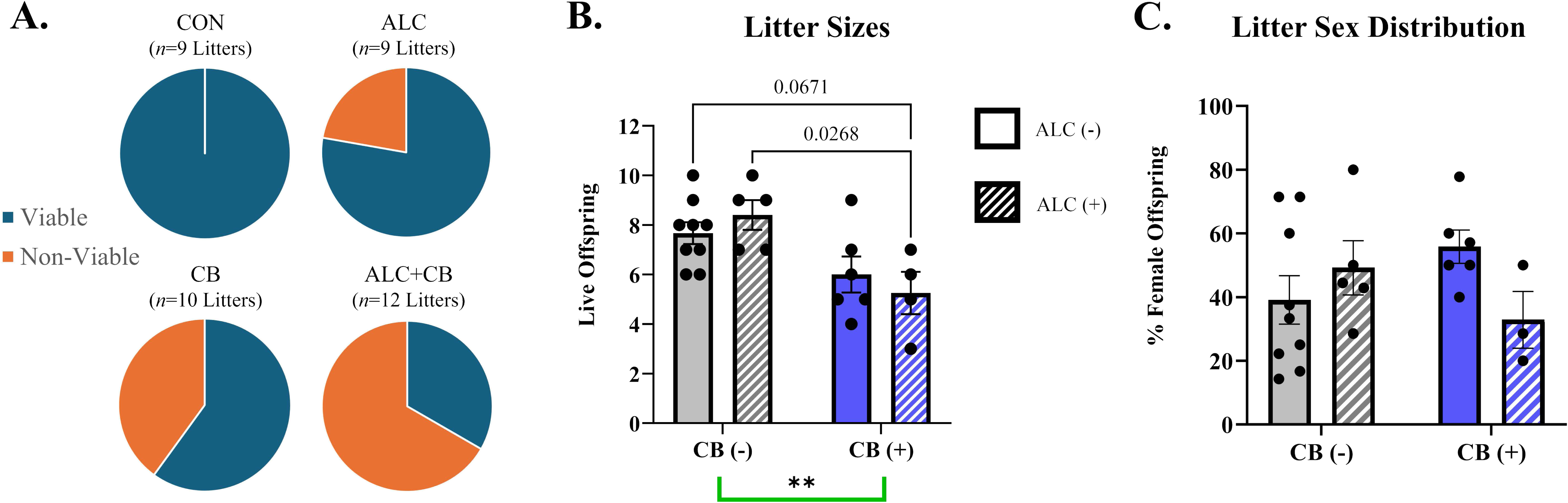
Litter characteristics at birth. **A)** Litter viability at birth differed between exposure groups, with ALC+CB offspring demonstrating the greatest loss of offspring viability compared to all other groups. **B)** Litter sizes among viable offspring demonstrate that ALC+CB litters produced fewer live offspring compared to CON and ALC exposures. *P-*values between pairwise comparisons are listed in graph. Offspring death between P0-P2 prevented quantification of two ALC litters. **C)** Litter sex distribution, assessed on Postnatal Day 2, revealed no significant differences in litter sex distribution as a result of exposure. Offspring death between P0-P2 prevented sex quantification among one ALC+CB litter. *Abbreviations:* CON = Control, ALC = Alcohol, CB = Cannabinoid, ALC + CB = Alcohol + Cannabinoid. **Symbols:** ** indicates *p* < 0.01

### Non-viable offspring demonstrate physical malformations, including impaired craniofacial development and inappropriate limb and organ development

Deceased offspring from six CB and ALC+CB litters exhibited distinct physical and developmental abnormalities (Table 2), with the severity of observable deficits recorded as deviation from age-matched, appropriately developed fetuses (scored as 0; Figure 2A). An ICC analysis was conducted to assess the inter-rater reliability of the ratings across four independent observers. Using a two-way random-effects model for absolute agreement with average measures, the ICC for observed physical deficits was 0.951, 95% CI [0.932, 0.965], and the ICC for estimated Theiler stage was 0.868, 95% CI [0.789, 0.922], indicating good-to-excellent reliability in assessments of deceased fetal metrics (Koo & Li, 2016).

**Figure 2.**
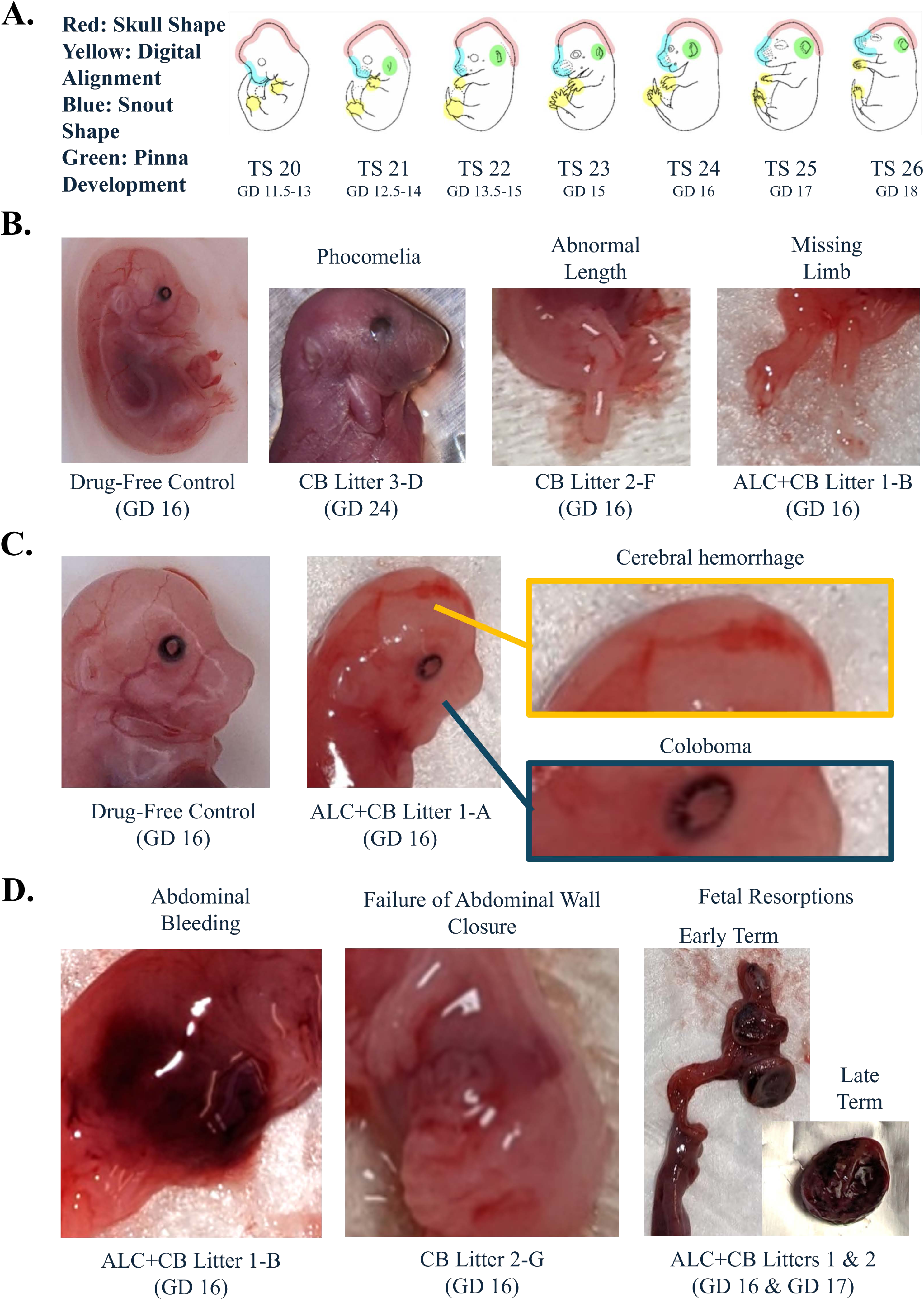
Characterization of non-viable offspring follow prenatal exposure. **A)** Theiler stages and associated Gestational Day equivalents, adapted from the EMAP eMouse Atlas (Richardson et al., 2013). Color-shaded regions indicate areas evaluated for developmental delays by researchers. **B)** Examples of limb deficits observed among non-viable offspring, including their associated exposure and litter ID #, alongside a drug-free control fetus. **C)** Example of an age-matched control fetus alongside a non-viable fetus with a blunted snout, cerebral hemorrhage (orange box) and coloboma (blue box). **D)** Examples of abdominal issues observed among non-viable offspring, including bleeding and failure for the abdominal wall to close, as evidenced by visible internal organs. *Abbreviations:* TS = Theiler stage, GD: Gestational Day, CB = Cannabinoid, ALC + CB = Alcohol + Cannabinoid. **Symbols:** ** indicates *p* < 0.01

**Table 2.**
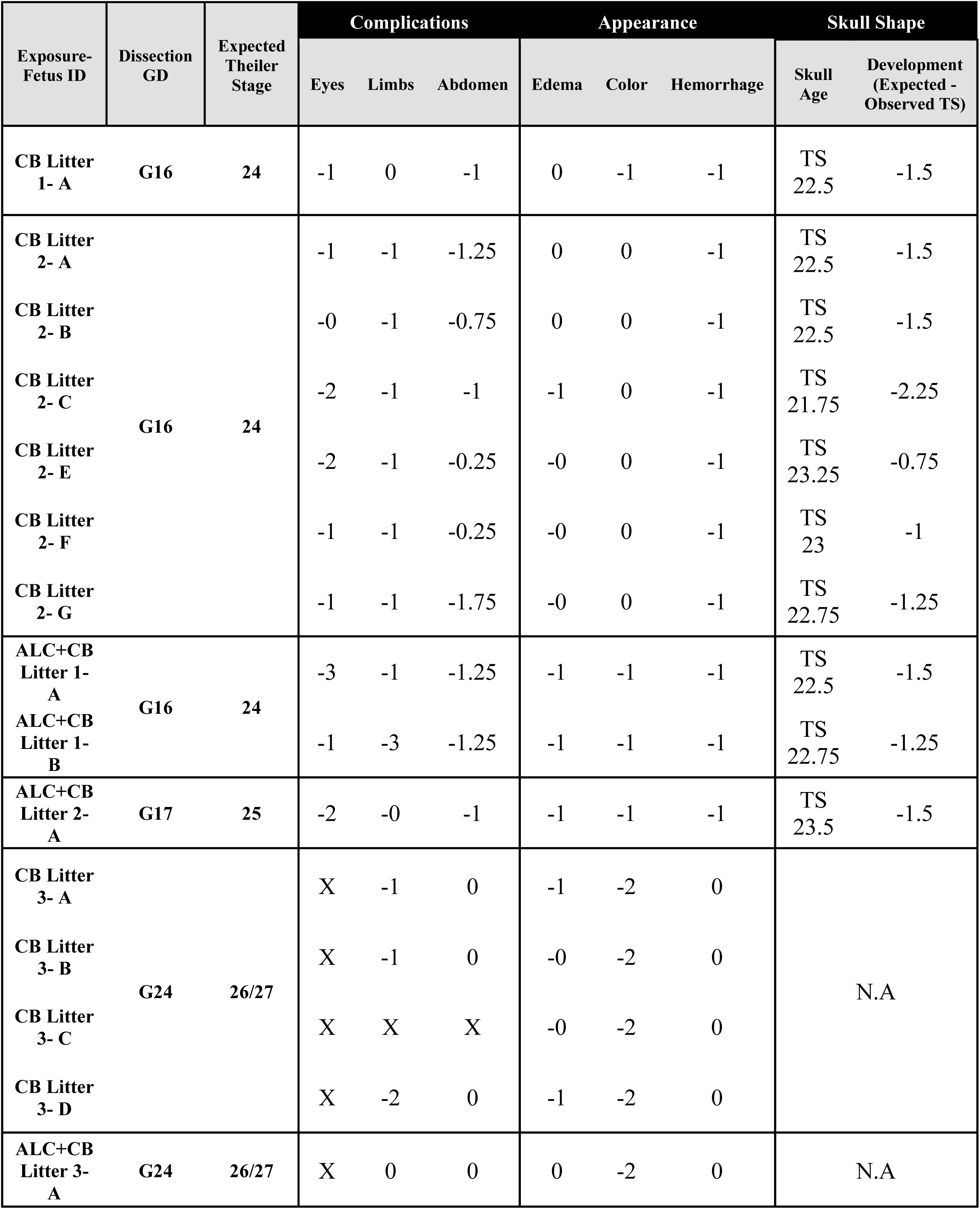
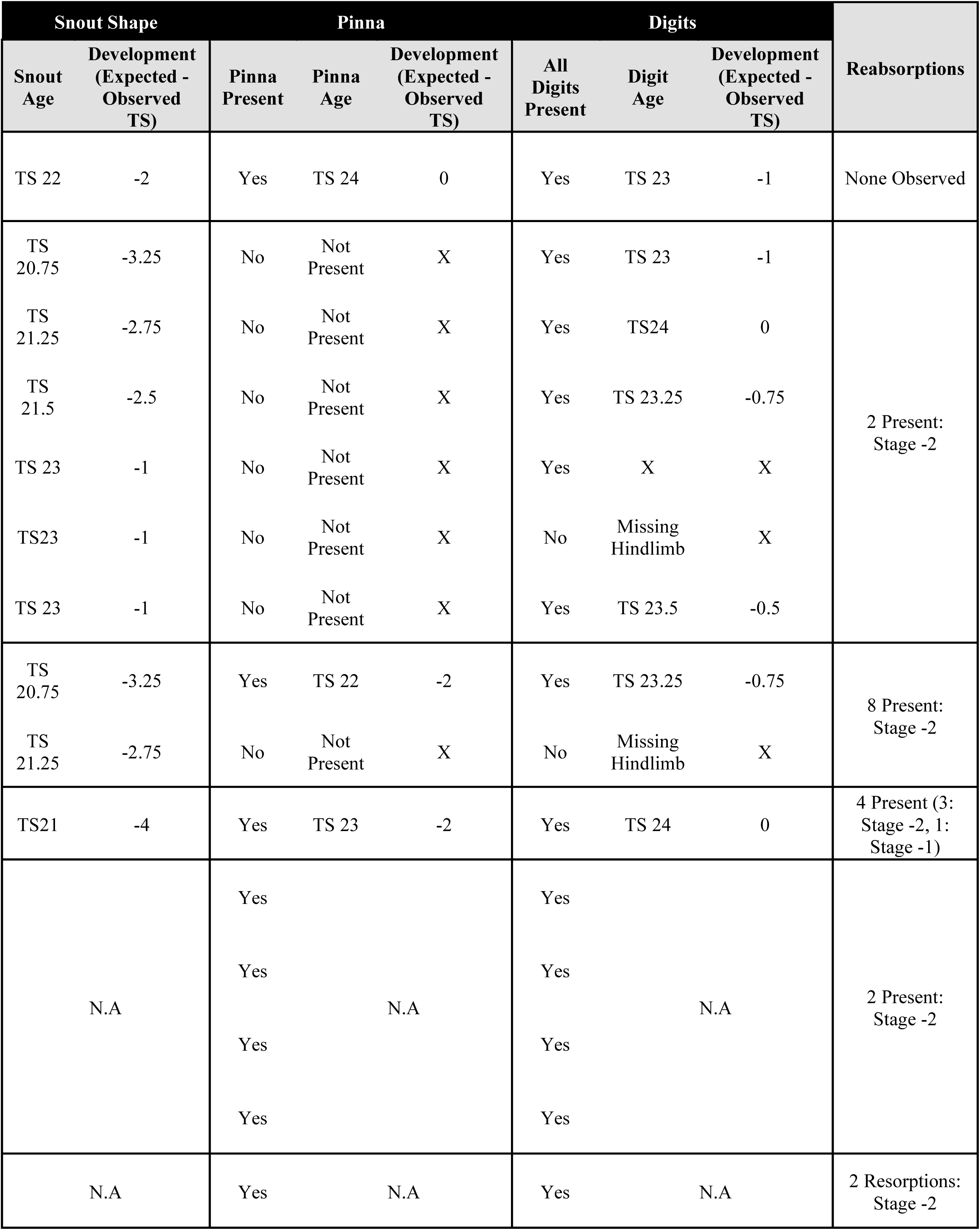
Average scores from four evaluators on offspring physical impairments and developmental delays in non-viable offspring from six litters.

Offspring from ALC+CB litters demonstrated greater overall severity in physical abnormalities compared to CB litters. In ALC+CB litters, iris abnormalities ranged from moderate non-circular shapes to severe colobomas (Figure 2C), with complication scores spanning -0.75 to -3, while CB litters showed milder iris defects, with scores ranging from -0.25 to -1.75. Limb abnormalities followed a similar trend, with ALC+CB litters displaying abnormal limb length and phocomelia, while CB-only litters were primarily affected by mild shortening/asymmetry (Figure 2B). Abdominal wall deficits, including visible organ exposure and bleeding, were more consistent in ALC+CB litters (-1.00 to -1.25), whereas CB litters exhibited a broader range of scores, including cases closer to normal development (-0.25 to - 1.75; Figure 2D). ALC+CB litters also displayed higher frequencies of fetal resorptions, with up to eight resorptions in advanced stages (-2), compared to an average of <2 resorptions in CB-only litters.

ALC+CB litters consistently exhibited skin discoloration, ranging from dark red to blue/purple (“-2” color scores). Notably, litters born after their anticipated delivery date demonstrated the most severe discoloration, potentially indicative of prolonged hypoxic distress. Edema was also more prevalent in ALC+CB litters, with ¾ offspring demonstrating at least moderate levels of swelling, while CB offspring occasionally exhibited no edema. Hemorrhagic features were prominent in both groups and exclusive to offspring born prior to their anticipated delivery date. Developmental delays were also prominent among offspring assessed prior to their anticipated delivery date. Skull development scores were relatively consistent between CB and ALC+CB offspring, with all premature litters demonstrating an average development delay of ∼1.5 Theiler stages. Notably, snout development was disrupted more by ALC+CB exposure, with offspring displaying underdeveloped snouts by ∼3 Theiler stages, while CB offspring averaged a lag of ∼2 Theiler stages. Pinnas were undetectable among ∼50% of all assessed offspring, independent of exposure, although this absence was exclusive to fetuses born prematurely. Finally, there were select cases of underdeveloped digit formation following either exposure, including two separate cases of incomplete hindlimb formation. All offspring assessed following their anticipated delivery date demonstrated appropriate pinna and digit development.

### Prenatal drug exposure contributes to increased early-life offspring mortality

In addition to periconceptual offspring mortality, postnatal offspring death was also observed only in drug-exposed litters. In two ALC litters and one ALC+CB litter, all offspring were cannibalized between PD0-2, which also prevented quantification of sex ratios, which were determined at PD2. After PD2, two CB litters demonstrated early neonatal mortality. From one litter, one male offspring died between PD2-4, while in a separate litter, one male offspring died between PD2-21 and two female offspring died between PD21-25. Control litters experienced no offspring mortality.

### Juvenile weight gains following prenatal polysubstance exposure are not sustained into adulthood

Sex-segregated offspring weights were first collected at weaning on PD21, and subsequently in adolescence (PD40) and young adulthood (PD80), prior to behavioral testing. Litter weights were averaged among male and female offspring, respectively, per litter. At PD21, there were statistically significant interactions for ALC x CB exposure for both male offspring [*F*(1, 18) = 10.85, *p* = 0.004] and female offspring [*F*(1, 19) = 7.576, *p* = 0.013], with ALC+CB offspring demonstrating the highest average juvenile weights among all exposure groups (Supplementary Tables 4&5; Supplementary Figure 1). Notably, ALC+CB offspring did not sustain their increased weights into PD40 or PD80, indicating that initial weight gain was not maintained during the transition into adolescence.

### Males and CB-exposed offspring require more trials to learn a Rotarod motor coordination task

During the first day of Rotarod testing, male and female mice were trained to perform the Rotarod task during a Habituation day, in which offspring were tasked with balancing on the Rotarod for 120 seconds at a speed of 5 rotations-per-minute (RPM). Animals who failed to balance for this long on the first trial were given two more opportunities to meet criteria, separated by a ten-minute inter-trial rest period. The number of trials required to meet the 120sec threshold were recorded, and only animals who met this threshold advanced to testing days. Out of all offspring tested, only one (ALC female) failed to meet the threshold; this subject was removed from further testing. On Habituation Day, both sex [*F*(1, 97) = 6.385, *p* = 0.013] and CB exposure [*F*(1, 97) = 4.720, *p* = 0.032], but not ALC exposure, affected the number of trials required to learn the Rotarod task: specifically, female offspring habituated to the task faster than males, independent of exposure, while CB-exposed offspring required more trials overall to learn the task than non-CB-exposed offspring (Supplementary Table 6; Figure 3A).

**Figure 3.**
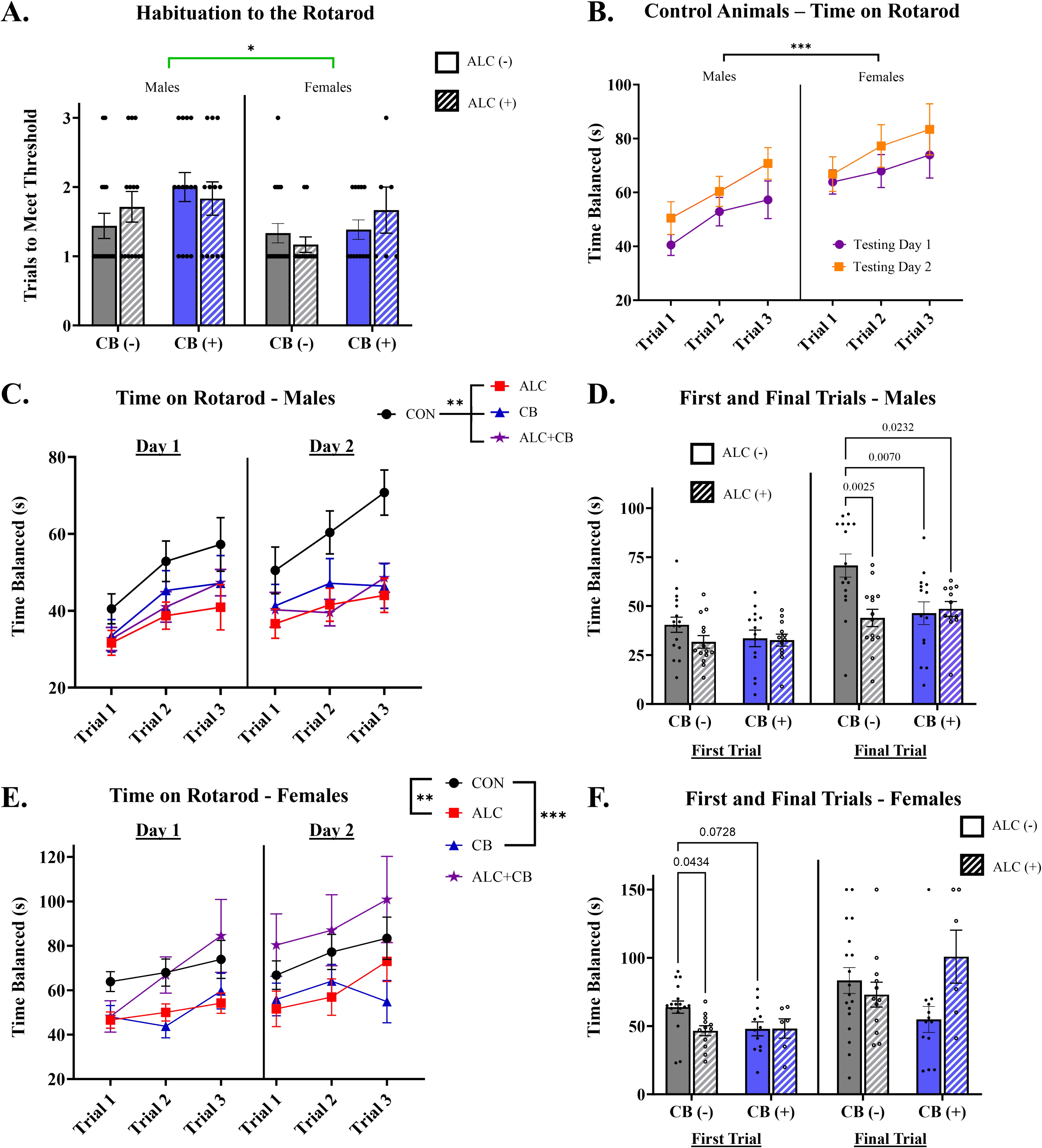
The effects of prenatal exposure on offspring rotarod coordination. **A)** Males and cannabinoid-exposed offspring required more trials to meet the threshold criteria for rotarod testing (120s balanced on the rod at 5 rotations/minute). **B)** Among control offspring, males and females improved in time balanced on the rod within and between testing days; however, females demonstrated the greatest overall balance. **C)** Among males, all forms of drug exposure reduced time spent balancing on the rotarod compared to controls. **D)** CON males do not differ from all other exposure groups in average time balanced on the rod during their first trial. However, by the final trial, CON males balance significantly longer than all other groups, indicating greater improvement in coordination across trials. *P-*values between pairwise comparisons are listed in graph. **E)** Among females, single-drug exposure, but not ALC+CB exposure, significantly reduced coordination on the rotarod. There were no differences between CON and ALC+CB females. **F)** During the first trial, CON females demonstrate better overall coordination than ALC and CB offspring. However, these differences are no longer significant by the final trial, indicating that female offspring improve in their coordination across trials. *P-*values between pairwise comparisons are listed in graph. *Abbreviations:* CON = Control, ALC = Alcohol, CB = Cannabinoid, ALC + CB = Alcohol + Cannabinoid. **Symbols:** * indicates *p* < 0.05, ** indicates *p* < 0.01, *** indicates *p* < 0.001

### In control animals, females demonstrate consistently longer balance on the Rotarod than males

When progressing onto Rotarod testing days, we uncovered further sex differences in the ability of control animals to balance (Figure 3B). Among drug-free controls, there were significant main effects of sex [*F*(1, 96) = 15.17, *p* < 0.001], trial number [*F*(2, 96) = 4.297, *p* = 0.016], and testing day [*F*(1, 96) = 6.672, *p* = 0.011] on rotarod performance (Supplementary Table 7). While both sexes demonstrated significant improvement in coordination on the rotarod within and between testing days, female offspring demonstrated better overall balance than males across six total trials.

### In male offspring, all prenatal exposures reduce coordination across Rotarod testing days

Across testing days and trials, assessments of Rotarod performance were compared between a) control offspring and each group of drug-exposed offspring, and b) single-drug exposed offspring and polysubstance-exposed offspring. In male offspring, all groups of drug-exposed offspring demonstrated significantly poorer coordination on the Rotarod than controls, as measured by time balanced on the rod and indicated by significant main effects of exposure: ALC: [*F*(1, 84) = 21.340, *p* < 0.001], CB [*F*(1,84) = 7.946, *p* = 0.006], and ALC+CB [*F*(1, 78) = 14.180, *p* < 0.001] (Figure 3C). However, offspring rotarod balance was not further impaired by polysubstance exposure compared to single-drug exposure: ALC [*F*(1,72) = 0.890, *p* = 0.349] or CB [*F*(1,72) = 0.246, *p* = 0.621]. Notably, within-group, only control males demonstrate significant main effects of ‘Trial’ [*F*(2,45) = 3.613, *p* = 0.035] and ‘Testing Day’ [*F*(1,45) = 9.553, *p* = 0.003], indicating consistent improvement in Rotarod coordination within each testing day and between testing days (Supplementary Table 8). In contrast, there were no significant main effects of ‘Trial’ and ‘Testing Day’ in any drug-exposed group (all *p*’s > 0.05; Supplementary Tables 9-11), indicating that prenatal drug exposure prevented improvement of performance on the rotarod across trials, both within and between testing days.

To assess the effects of ALC and CB exposures on offspring rotarod performance in males, the first and final trials of the Rotarod task were assessed for interactions between exposure group and trial (Figure 3D). There was a significant main effect of ALC exposure [*F*(1,52) = 5.105, *p* = 0.028] on Rotarod coordination, a significant interaction between ALC x CB exposure [*F*(1,52) = 5.919, *p* = 0.019], and a significant three-way interaction between CB exposure x ALC exposure x Trial [*F*(1,52) = 4.362, *p* = 0.042] (Supplementary Table 12). During the first trial, post-hoc analyses revealed that there were no significant differences in time balanced on the rod, between control offspring and all exposure groups (all *p*’s > 0.05; Supplementary Table 13). However, on the final trial, control offspring balanced significantly longer than ALC offspring (*p*=0.003), CB offspring (*p*=0.007), and ALC+CB offspring (*p*=0.023). There were no significant differences between single-drug exposed offspring and polysubstance-exposed offspring during the first or last trials (all *p*’s > 0.05; Supplementary Table 13).

### In female offspring, prenatal single-drug exposure, but not polysubstance-exposure, reduces coordination on a Rotarod task

In female offspring, ALC exposure [*F*(1, 84) = 12.850, *p* < 0.001] and CB exposure [*F*(1, 87) = 13.100, *p* < 0.001], individually, reduced offspring coordination on the Rotarod compared to controls (Figure 3E). However, Rotarod balance was not significantly affected by polysubstance exposure when compared to control offspring [*F*(1, 66) = 0.601, *p* = 0.441]. In contrast, time balanced on the Rotarod was significantly longer in ALC+CB female offspring when compared to ALC [*F*(1, 48) = 10.590, *p* = 0.002] and CB [*F*(1, 51) = 9.010, *p* =0.004] offspring. Within-group, no exposure produced a significant main effect of ‘Trial’ (all *p*’s > 0.05; Supplementary Tables 14-17), indicating that performance within-day was relatively consistent for all groups. There was a significant main effect of ‘Testing Day’ in ALC [*F*(1, 33) = 5.499, *p* = 0.025] and ALC+CB offspring [*F*(1, 15) = 16.350, *p* = 0.001], with offspring improving in their rotarod coordination between Days 1 and Day 2 of Testing. In contrast, there was no significant effect of ‘Testing Day’ in CON [*F*(1, 50) = 1.586, *p* = 0.214] or CB offspring [*F*(1, 34) = 1.754, *p* = 0.194], indicating no significant improvement in rotarod balance between testing days.

As with male offspring, the first and final trials of the Rotarod task were assessed for interactions between exposure group and ‘Trial’ in female offspring (Figure 3F). There was a significant interaction between ALC x CB exposure [*F*(1,45) = 5.220, *p* = 0.027], a significant interaction between ALC exposure x ‘Trial’ [*F*(1,43) = 6.996, *p* = 0.011], and a significant three-way interaction between CB exposure x ALC exposure x ‘Trial’ [*F*(1,43) = 4.301, *p* = 0.044] (Supplementary Table 18). During the first trial, post-hoc analyses revealed a significant reduction in time balanced on the Rotarod in ALC offspring (*p*=0.043) and a non-significant reduction in CB offspring (*p*=0.073), with no effect on performance in ALC+CB offspring (*p*=0.215). In contrast, there were no significant differences in time balanced on the rotarod between any exposure groups during the final trial (all *p*’s > 0.05; Supplementary Table 19).

### Prenatal polysubstance exposure decreases time exploring the center of the open field, while increasing hyperactivity measures

Offspring were next assessed in an open field assay, in which subjects could freely explore the apparatus for ten minutes, with their activity recorded for sex and exposure-induced changes in exploratory behaviors. Foremost, control males and females were compared for innate sex differences in open field activity. Control male offspring, on average, spent more time exploring the center of the open field arena compared to control female offspring [*t*(1, 32) = 2.827, *p* = 0.008] (Figure 4A). However, sex did not impact additional open field measures in control offspring, including times entering the center, distance travelled throughout the open field arena, and average speed travelled (Supplementary Figure 2; Supplementary Table 20).

**Figure 4.**
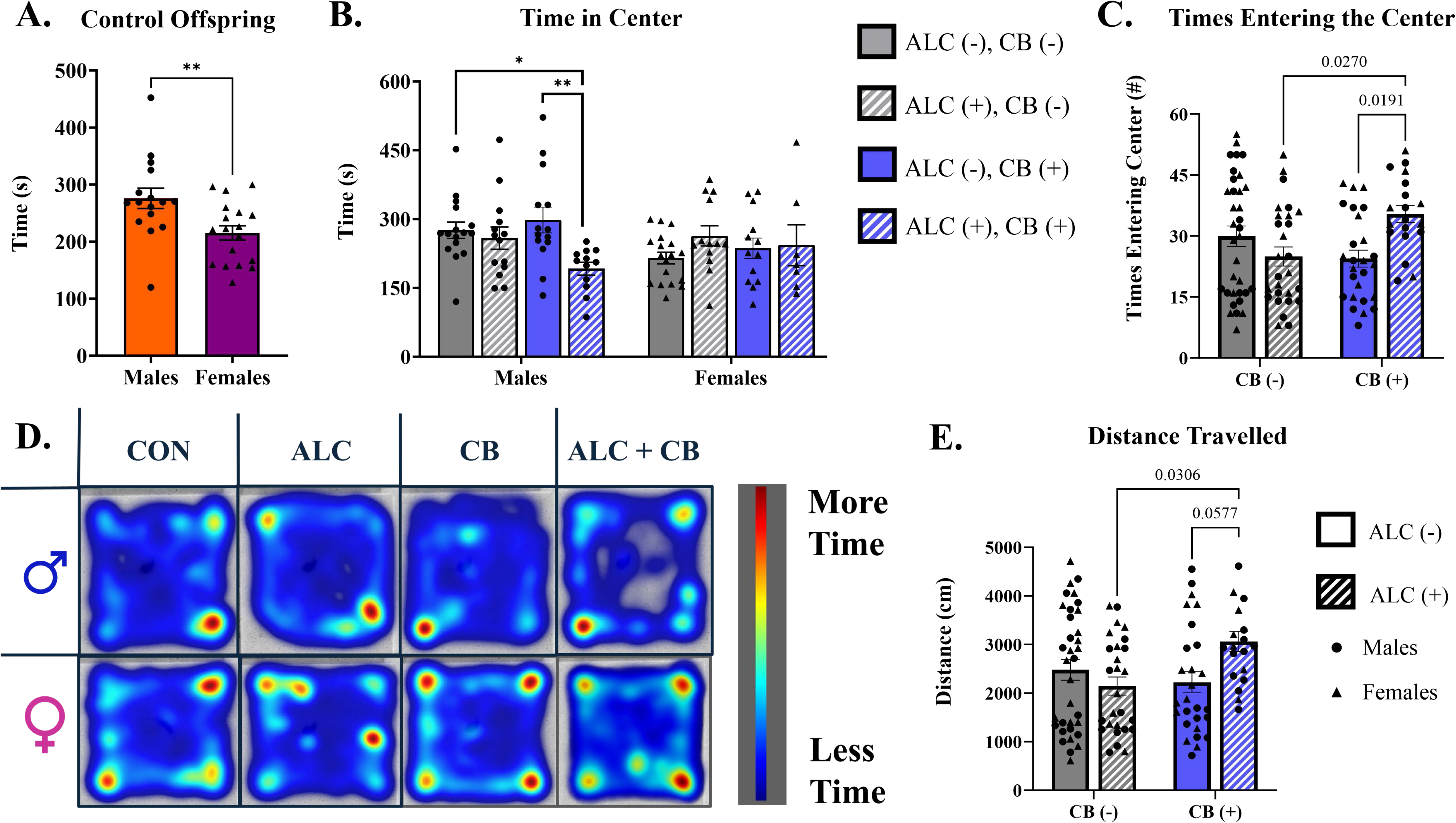
The effects of prenatal exposure on offspring open field measures. **A)** Among control offspring, males spent more time in the center of the open field arena than females. **B)** Exclusively in males, ALC+CB exposure reduces time spent in the center of the open field arena compared to CON and CB exposures. There is no effect of exposure in females. **C)** ALC+CB exposure increases the number of entries into the open field arena compared to single-drug exposures. *P-*values between pairwise comparisons are listed in graph. **D)** Heatmaps representing average activity patterns among exposure groups and sexes during the open field test. E) ALC+CB exposure increases the distance travelled throughout the the open field arena compared to single-drug exposures. *P-*values between pairwise comparisons are listed in graph. *Abbreviations:* CON = Control, ALC = Alcohol, CB = Cannabinoid, ALC + CB = Alcohol + Cannabinoid. **Symbols:** * indicates *p* < 0.05, ** indicates *p* < 0.01

When assessing time spent in the center of the open field across exposures, there was a significant interaction between CB x ALC Exposure [*F*(1,55) = 4.136, *p* = 0.044] and Sex x ALC Exposure [*F*(1,99) = 7.725, *p* = 0.007] (Figure 4B). In post-hoc assessments, ALC+CB male offspring spent significantly less time in the center of the open field compared to control (*p*=0.048) and CB males (*p*=0.010) (Supplementary Table 21), whereas no effect of exposure was observed in any female offspring (Supplementary Table 22; Figure 4D). In assessments of entries into the center of the open field, there was a significant interaction between CB x ALC Exposure [*F*(1,98) = 10.590, *p* = 0.002], with no influence of sex as a main effect or interacting variable (Supplementary Table 23). Data were subsequently collapsed by sex to perform post-hoc comparisons of exposure groups. ALC+CB offspring demonstrated significantly more entries into the center of the open field compared to ALC offspring (*p*=0.027) and CB offspring (*p*=0.019), but not control offspring (*p*=0.406) (Supplementary Table 24; Figure 4C).

Two measures within the open field were assessed as metrics of hyperactivity: average speed travelled during the testing session, and total distance travelled for the length of the session. For average speed, there was a statistically significant interaction between ALC x CB Exposure [*F*(1,99) = 7.540, *p* = 0.007], and no effect of sex as a main effect or interacting variable (Supplementary Table 25). This finding was similarly observed in distance travelled, with only a significant interaction between CB x ALC Exposure [*F*(1,77) = 7.544, *p* = 0.007] (Supplementary Table 26). Data were subsequently collapsed by sex to perform post-hoc comparisons of exposure groups. ALC+CB offspring demonstrated higher average speeds while traveling through the open field compared to ALC offspring (*p*=0.031) and CB offspring (*p*=0.058), but not control offspring (*p*=0.255) (Supplementary Table 27). Similarly, ALC+CB offspring travelled farther distances during the open field test than ALC offspring (*p*=0.031) and CB offspring (*p*=0.058), but not control offspring (*p*=0.254) (Supplementary Table 28; Figure 4E).

## Discussion

Recent studies in human populations have associated self-reported prenatal cannabis exposure with increased risk of offspring mortality (Gretchen Bandoli et al., 2023; Crosland et al., 2024; Shi et al., 2021; Varner et al., 2014), pre-term delivery (Crosland et al., 2024; Luke et al., 2022; Petrangelo et al., 2019; Shi et al., 2021), and Sudden Infant Death Syndrome (SIDS; (Scragg et al., 2001)). Our data indicate that the link with perinatal mortality and preterm birth is likely to be causal. Importantly, we observed elevated mortality through the pre-pubertal period, an outcome that needs to be assessed in human populations as well. We also found that the impact of prenatal cannabinoid exposure on offspring mortality was further exacerbated by simultaneous alcohol exposure, resulting in fewer live births and reduced litter sizes. This is significant, since polysubstance use in the context of risky alcohol consumption is increasingly common. There are a few pre-clinical studies on the combined effects of alcohol and cannabinoids, and our results are largely consistent with those studies. For example, an early investigation showed that combined prenatal alcohol (2 g/kg) and Δ9-tetrahydrocannabinol (THC; 100 mg/kg) exposure from GD 1-15 resulted in significant increases in fetotoxicity among mice and rats compared to single-drug-exposed and drug-free offspring (Abel, 1985).

More recently, Breit and colleagues (2019a) demonstrated greater lifetime mortality in offspring following co-exposure to alcohol (5.25 g/kg/day) and cannabinoids (CP55940; 0.4 mg/kg/day) from PD 4-9 compared to single-drug-exposed and drug-free offspring. Collectively, these and our study show that prenatal co-exposure to alcohol and cannabinoids may confer additional risks for delivery complications and offspring mortality, independent of the developmental window for exposure (prenatal vs early postnatal) or type of cannabinoid compound (naturally occurring vs synthetic). Notably, alcohol exposure alone in our model did not significantly affect offspring viability or litter size. While some clinical studies have linked prenatal alcohol exposure to pre-term delivery and SIDS (Beth A Bailey & Robert J Sokol, 2011), others report no association between alcohol exposure and delivery complications, indicating that these effects may depend on the dose and timing of exposure (Lundsberg et al., 2015; Pfinder et al., 2013).

Prenatal exposure to alcohol and cannabinoids, both individually and in combination, can also result in significant growth deficits and adverse physical development. Prenatal alcohol exposure is well-documented to cause persistent growth deficits, affecting offspring weight, height, and head circumference, with delays extending into adolescence (Carter et al., 2013; Day et al., 1989; Day et al., 1994; Pielage et al., 2023). These outcomes, while dose-dependent, can occur following even low levels of alcohol consumption during pregnancy (Day et al., 2002).

Similarly, prenatal cannabinoid exposure has been associated with growth restrictions and an increased likelihood of being born small for gestational age (Crosland et al., 2024; Huizink, 2014; Natale et al., 2020). In co-exposure models, more severe growth deficits have sometimes been observed in co-exposed offspring compared to single-drug-exposed offspring (Breit et al., 2019a, 2019b). However, other studies have not replicated these findings (Breit et al., 2020), indicating variability in outcomes. Interestingly, in our study, no growth restrictions were observed in viable offspring subjected to prenatal substance exposure. Instead, ALC+CB offspring demonstrated initial weight gains by PD 21. These weight increases may reflect the smaller litter sizes in ALC+CB groups, which could potentially increase access to maternal care and nutrition among surviving offspring. However, this early advantage did not persist; by PD 40 and PD 80, the weights of ALC+CB offspring were comparable to those of all other exposure groups.

A lack of weight differences among drug-exposed offspring may reflect a survival bias, as no offspring born prematurely survived to term. Underdeveloped animals may have died, skewing our results to only the most resilient litters. This possibility is supported by our physical examinations of non-viable offspring, with consistent physical malformations observed across offspring from six cannabinoid or dual-exposed litters, revealing hypodevelopment of the face, abdomen, and limbs. Independently, prenatal alcohol exposure is well-documented to produce craniofacial dysmorphologies characteristic of Fetal Alcohol Spectrum Disorder (FASD). These anomalies include alterations in the midface, nose, lips, and eyes, often resulting in a general recession of the midface and superior displacement of the nose (Muggli et al., 2017; Suttie et al., 2013). Studies using rodent models have further demonstrated that alcohol exposure during critical periods of development can produce significant craniofacial malformations. [see review: (Petrelli et al., 2018)]. Prenatal cannabinoid exposure has similarly contributed to the development of microphthalmia, iridial colobomas, facial clefts, and holoprosencephaly (Gilbert et al., 2016), as well as limb reduction, musculoskeletal deficiencies, and cardiovascular issues (Reece & Hulse, 2019). Moreover, limited studies on co-exposure suggest synergistic effects, such as enhanced craniofacial defects mediated by disruptions in cannabinoid receptor 1 (CNR1) signaling and the Sonic Hedgehog pathway (Boa-Amponsem et al., 2019; Fish et al., 2019).

Unfortunately, as our morphometric analyses were limited to litters with complete mortality, that were predominantly those exposed to cannabinoids, we could not evaluate whether combined exposure exacerbates outcomes relative to single-drug exposure in this cohort.

Notably, non-viable cannabinoid-exposed offspring exhibited evidence of intra-abdominal hemorrhage and preterm fetuses exhibited failure of abdominal wall closure. One potential cause is prenatal inflammation, which is strongly associated with preterm birth and fetal hemorrhaging. For example, lipopolysaccharide (LPS) exposure *in utero* induced liver hemorrhaging in ∼70% of fetuses, while control fetuses remained unaffected (Ernst et al., 2010). Alternatively, exposure to alcohol and cannabinoids may dysregulate the mechanistic target of rapamycin (mTOR) signaling pathway, which governs vertebrate growth, embryonic development, and organogenesis (Hwang et al., 2008). Acute exposure to both alcohol and THC can increase mTORC1 signaling (Neasta et al., 2014), and uterine mTOR hyperactivation has been implicated in preterm birth in mice (Hirota et al., 2011). While prenatal alcohol exposure has been shown to alter mTOR activity in the fetal hippocampus (Lee et al., 2020), cerebellum, and skeletal muscle (Sawant et al., 2020), the potential role of mTOR dysregulation in drug-induced uterine or systemic fetal outcomes warrants further investigation. Importantly, the relationship between intrauterine inflammation and compromised fetal health may be mediated by mTOR signaling, as LPS administration has been associated with acute elevations in mTOR signaling in the fetal brain (Dong et al., 2020).

Additional mechanisms for increased fetal mortality and abdominal bleeding include deficiencies in blood coagulation factors. Deficiencies in blood coagulation Factor X (Dewerchin et al., 2000) and prothrombin clotting factor 2 (Xue et al., 1998) have been associated with embryonic lethality and fatal neonatal bleeding, while Factor XIII A subunit-deficient mice experience severe uterine bleeding and spontaneous miscarriage (Koseki-Kuno et al., 2003).

However, the effects of prenatal alcohol and cannabinoid exposure on coagulation factors are currently poorly understood. Finally, fetal hypoxia and compromised blood flow/ischemia can contribute to vascular stress and breakdown, leading to internal bleeding. We recently demonstrated that both alcohol and cannabinoid exposures reduce fetal cerebral blood flow *in utero* (Rouzer et al., 2024), with combined exposure augmenting blood flow deficits through the internal carotid artery. These reductions corresponded with instances of intrauterine growth restriction, emphasizing the need for further research to determine whether recovering losses in fetal blood flow could mitigate embryonic lethality and developmental malformations associated with prenatal drug exposure.

Significant sex differences emerged when assessing coordination abilities in prenatally exposed young adult offspring. Male offspring prenatally exposed to cannabinoids required significantly more trials to habituate to the Rotarod task on the first day, aligning with prior findings that THC-exposed adolescent offspring required more trials to achieve their first success performance in a parallel bar test (Breit et al., 2022). Among control offspring, females consistently outperformed their male siblings, balancing longer on the Rotarod, although both sexes improved across testing trials. This observation is consistent with reports that B6 mouse strains exhibit motor learning through repeated testing (McFadyen et al., 2003), and B6J female mice can balance longer on the rotarod than male counterparts (Ashworth et al., 2015). Analysis of exposure effects within sexes revealed distinct patterns. In males, all prenatal exposure groups showed reduced balancing time on the Rotarod compared to unexposed controls, with the most pronounced deficits occurring in the final trial. In contrast, females exhibited impairments exclusively in single-drug exposure groups, with the most significant deficits observed during the first trial, suggesting that drug-exposed females displayed “catch-up” performance with repeated testing. These findings are consistent with previous studies on prenatal single-drug exposure, which report impairments in motor learning and coordination (Breit et al., 2019a; Breit et al., 2022; Connor et al., 2006; Reekes et al., 2016; Thomas et al., 2000). Interestingly, the deficits observed in ALC males resemble a study of six-month-old male mice, where moderate prenatal alcohol exposure did not affect initial performance but reduced improvement across testing days (Reekes et al., 2016). However, additional studies of prenatal alcohol or cannabinoid exposure report no significant coordination impairments following single-drug exposure (Breit et al., 2019a; Breit et al., 2022; Heaton et al., 2022).

These inconsistent outcomes in coordination deficits extend to preclinical investigations of ALC+CB exposure. For example, in a parallel bar test, co-exposure to CP-55940 and alcohol amplified adolescent (∼PD 30) coordination deficits compared to single-drug exposure, particularly in female offspring (Breit et al., 2019a). However, the same research group found that alcohol co-exposure did not exacerbate THC-induced deficits, with THC alone driving reduced coordination (Breit et al., 2022). In our study, co-exposure did not augment outcomes from single-drug exposure. Co-exposed offspring generally exhibited impairments comparable to single-drug exposed groups, except for ALC+CB females, who performed similarly to controls. Notably, ALC+CB females also had the lowest survival rate in our cohort (*n*=6 from 13 litters), suggesting their strong performance may reflect a survival bias favoring resilient individuals.

Variability in reported outcomes within this field of investigation likely arises from differences in the type of cannabinoid used, dosage, route of administration, and timing of gestational exposure, among other experimental variables. These findings underscore the importance of studying co-exposures alongside comparable single-drug exposures, accounting for experimental factors that influence behavioral outcomes.

In contrast to coordination assessments, sex differences were minimal in open field activity measures. In control offspring, males did spend more time in the center of the OF than females; otherwise, no sex differences were observed in open field measures. Furthermore, unlike coordination measures, single-drug exposures did not alter open field behaviors. However, ALC+CB exposure significantly reduced center time in males compared to controls and CB-only offspring. Notably, ALC+CB offspring entered the center more frequently than ALC or CB groups, but this likely reflects polysubstance-induced hyperactivity, as co-exposed offspring also exhibited significantly increased speed and distance travelled. Overall, ALC+CB offspring displayed a thigmotaxic behavioral phenotype, characterized by faster movement along the periphery of the open field.

The open field test is a validated tool for assessing hyperactivity and anxiety-like behaviors in rodents (Kraeuter et al., 2019). Hyperactivity and inattention are common symptoms expressed by individuals with prenatal exposure to alcohol (Dodge et al., 2023; Han et al., 2015; Infante et al., 2015) or cannabinoids [reviewed in (Ikeda et al., 2022)] throughout the lifespan. In the open field test, avoidance of, and reduced time spent in the center of the open field may reflect increased anxiety-like behavior. In rodents, avoidance of the center reflects increased anxiety-like behavior, aligning with the high prevalence of anxiety disorders in individuals with prenatal alcohol exposure (Burgess & Moritz, 2020; Hellemans et al., 2010), who may first demonstrate symptoms in early childhood (O’Connor & Paley, 2009). Similarly, prenatal cannabinoid exposure can also heighten the risk for offspring anxiety disorders (Grant et al., 2018; Nashed et al., 2021).

When reviewing existing preclinical studies, it’s noteworthy that assessments of effects of ALC exposure on open field activity have produced mixed results, with some researchers reporting no effects of prenatal exposure (Allan et al., 2003; Downing et al., 2008), while others report alcohol-induced hyperactivity in the open field (Randall et al., 1985), or hypoactivity (Rouzer et al., 2017; Schambra et al., 2016). Similarly, preclinical investigations of prenatal cannabinoid exposure have uncovered reduced time in the center of the open field (Cupo et al., 2024; Newsom & Kelly, 2008), and increased anxiety-like behaviors and attention deficits following THC exposure (Navarro et al., 2024). However, findings vary depending on the type and dose of cannabinoid, route of administration, rodent strain, and timing of exposure [as reviewed in (Higuera-Matas et al., 2015)]. Factors of offspring age and sex (Higuera-Matas et al., 2015; Osterlund Oltmanns et al., 2022; Schambra et al., 2016; Andrew F Scheyer et al., 2019) further mediate the effects of prenatal substance exposure on attention, hyperactivity, and anxiety-like behaviors.

It is notable that within our own study, individual exposures to alcohol or cannabinoids did not alter open field behaviors, but simultaneous prenatal exposure produced a phenotype of hyperactivity and anxiety-like behavior consistent with FASD. Our findings align with prior research demonstrating augmented hyperactivity following co-exposure. For instance, Ornelas et al (2024) reported increased adolescent hyperactivity levels in co-exposed males compared to cannabinoid-exposed-only offspring, at CP-55940 doses <50% lower than those used in our study. Separately, zebrafish co-exposed to ethanol and a cannabinoid receptor 1 agonist demonstrated significantly increased swimming activity, an effect that was absent following equivalent single-substance exposures (Boa-Amponsem et al., 2019). Similarly, third-trimester-equivalent co-exposure to alcohol and CP-55940 (0.4mg/dL) augmented locomotor activity beyond individual substance effects (Breit et al., 2019b). Collectively, these studies highlight hyperactivity as a robust phenotype of prenatal co-exposure, supporting the need for further research into the distinct and synergistic effects of co-exposures compared to single-drug exposures.

### Limitations

Our study employed a controlled model of prenatal alcohol and cannabinoid co-exposure to investigate offspring outcomes, building on prior assessments of how this exposure paradigm influences intrauterine fetal blood flow (Rouzer et al., 2024). As previously addressed, while this model provides valuable insights, it has several inherent limitations. First, our use of vaporized ethanol exposure from G12-15 allowed for precise control of maternal blood alcohol concentrations (Rouzer et al., 2024) and exposure duration across pregnancies.

This standardization minimizes variability, but may inadvertently introduce maternal stress during pregnancy, potentially confounding offspring outcomes. Similarly, cannabinoid exposure relied on intraperitoneal injections of the synthetic cannabinoid CP-55,940, rather than natural phytocannabinoids such as THC or CBD. While this method ensured consistent dosing and pharmacokinetics compared to inhalation or oral routes, it limits translational applicability. CP-55,940, a potent and selective agonist of CNR1 receptors, serves as a suitable proxy for synthetic cannabinoid exposure, but does not reflect the complex cannabinoid profiles of recreational or medicinal cannabis products. Future studies should explore a broader range of exposure methods, doses, and cannabinoid compositions to enhance the translational relevance of these findings.

A notable limitation of this model was the significant mortality observed in offspring following co-exposure to alcohol and cannabinoids. Mortality was disproportionately high in co-exposed litters, underscoring the severe teratogenic risks of combined exposure. However, this reduction in offspring survival reduced sample sizes for subsequent behavioral testing, possibly compromising statistical power. Moreover, physiological adaptations that contributed to offspring survival in the co-exposure group may have also introduced ‘survivor’ biases in behavioral performance. Future research should consider dose-response studies to identify exposure thresholds that significantly affect mortality, enabling better modeling of varied exposure levels. Finally, our analyses of physical malformations were conducted *post hoc* due to the marked mortality observed in cannabinoid-exposed and co-exposed groups. Non-viable offspring from control and alcohol-only groups were not included, limiting direct comparisons across all exposure conditions and potentially obscuring group-specific effects. To address this, future studies should incorporate planned, age-matched dissections and morphometric assessments across all experimental groups. This approach would enable more comprehensive evaluations of fetal abnormalities and their associations with different forms of prenatal drug exposure.

### Conclusions

In this study, we demonstrated that prenatal alcohol and synthetic cannabinoid co-exposure significantly compromised offspring viability, with higher rates of pre- and postnatal mortality compared to single-drug exposures or controls. Behavioral assessments in surviving offspring revealed distinct sex-specific effects on motor coordination and activity levels, with co-exposed males and females displaying differing patterns of impairment or resilience. Notably, co-exposure augmented hyperactivity and anxiety-like behaviors in offspring, consistent with phenotypes observed in human populations with prenatal drug exposure. Despite the wealth of knowledge regarding the effects of individual alcohol or cannabinoid exposures, polysubstance exposure remains underexplored, particularly in models mimicking real-world patterns of drug use during pregnancy. Further investigation is needed to elucidate the neurobiological mechanisms underlying these outcomes, including possible contributions from altered mTOR signaling, intrauterine inflammation, and vascular dysregulation, as well as neurotransmitter systems that are similarly targeted by individual exposures [as detailed in (Rouzer et al., 2023)]. Such research is essential for advancing our understanding of the long-term consequences of prenatal co-exposure, and developing targeted interventions to mitigate its effects on offspring health and development.

## Acknowledgements

The authors would like to thank Dr. Karienn Montgomery for providing training in the use of behavioral equipment. Furthermore, the authors would like to acknowledge the Neuroscience & Experimental Therapeutics Neurobehavioral Core for providing the space and resources to perform the behavioral assessments included in this manuscript.

## Contributions

SKR and RCM contributed to the conception/design of this research, and SKR was responsible for the acquisition and data curation. SKR, MD, AG, and AB contributed to the analysis of collected data, as well as the original manuscript preparation. All authors read and approved the final version of the manuscript.

## Funding Sources

This study was funded by grants from the National Institute of Alcohol Abuse and Alcoholism: F32 AA029866 (SKR), L40 AA030427 (SKR) & R01 AA028406 (RCM).

---

**Supplementary Figure 1.**
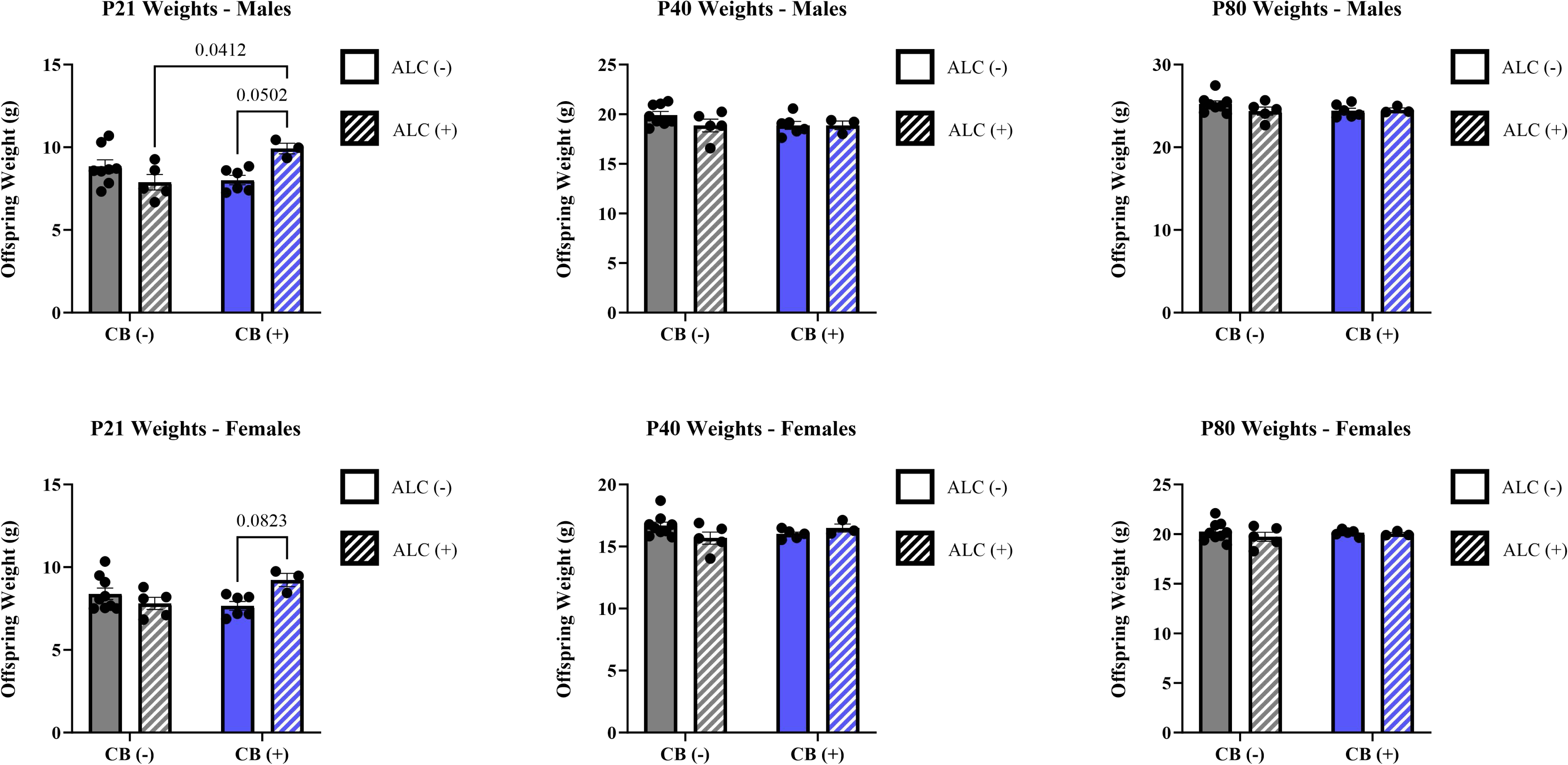
Early life weights (g) across exposure groups. Although males and females with ALC+CB exposure initially demonstrate minor increases in weight on Postnatal Day 21, these increases are not sustained into adolescence or young adulthood.

**Supplementary Figure 2.**
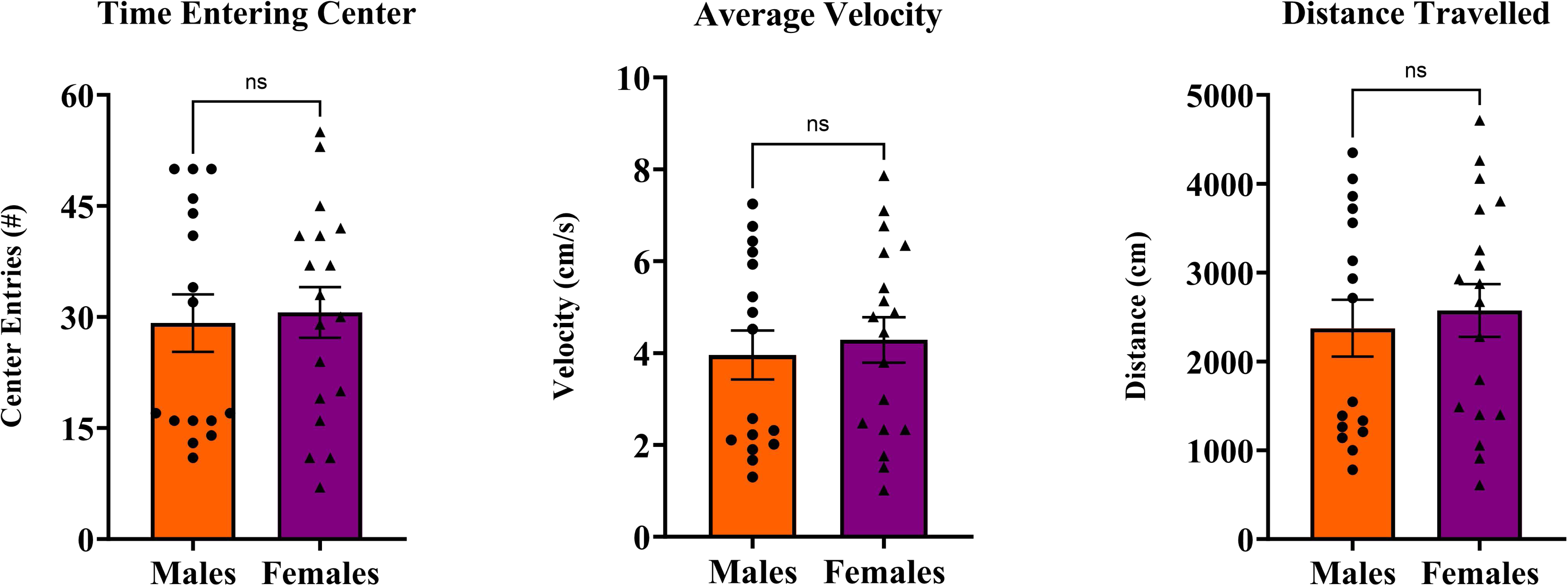
Open field measures among control offspring. Males and females do not differ in their entries into the center of the open field arena, nor their average velocity or distance travelled during the open field test. *Abbreviations:* NS = not statistically significant

## Notes

### Competing Interest Statement

The authors have declared no competing interest.

